# Polar auxin transport regulates initial auxin patterning and polar nuclear migration during lateral root initiation

**DOI:** 10.64898/2026.09.04.749482

**Authors:** Sanae Kaneta, Tatsuo Kakimoto

## Abstract

In *Arabidopsis thaliana*, local auxin accumulation in xylem-pole pericycle (XPP) cells specifies lateral root founder cells, which establish *de novo* cell polarity manifested by polar nuclear migration and subsequent asymmetric division. Here, we developed a fast-maturing, XPP/early lateral root primordium (LRP)-specific R2D2 auxin reporter (fx-R2D2) to visualize auxin dynamics with high spatiotemporal resolution during LRP initiation. We demonstrate that auxin increases broadly on the convex side of XPP during root bending. Within this region, cells showing particularly strong auxin responses emerged, and subsequently underwent polar nuclear migration and asymmetric division, initiating an LRP. When two LRPs initiated in close proximity, auxin levels in one rapidly decreased after the cell division, followed by cessation of further division. These findings provide a clear picture of auxin dynamics during LRP initiation. Under uniform auxin treatment, *de novo* LRP formation events occur. Auxin treatment initially induced a uniform auxin response at a high level, followed by localized auxin depletion in regions where LRPs did not subsequently form, demonstrating that auxin depletion as well as accumulation shapes the auxin patterns during LRP initiation. Furthermore, we reveal that local PIN-mediated polar auxin transport regulates both *de novo* auxin patterning and polar nuclear migration during LRP initiation.

**Highlight:** A novel high-resolution tissue-specific auxin reporter and quantification of nuclear migration reveal the role of PIN-mediated auxin transport in initial auxin patterning and polar nuclear migration during lateral root initiation.

## Introduction

Auxin plays an integral role in the formation of new organs in plant development. Local auxin maxima, which are typically created by self-organization through polar auxin transport within the tissue, trigger the formation of organs or tissue patterns including embryogenesis (Benková et al., 2003), leaf primordium formation (Okada et al., 1991; Gälweiler et al., 1998; Reinhardt et al., 2003; Jönsson et al., 2006), vascular tissue patterning (De Rybel et al., 2014), lateral root formation (Dubrovsky et al., 2008), and tissue regeneration (Efroni et al., 2016). These events involve cellular processes such as cell division, cell differentiation and cell polarity establishment. In *Arabidopsis thaliana*, lateral roots (LRs) are derived from the pericycle cells adjacent to the xylem (xylem-pole pericycle, XPP). Auxin accumulation in one or two longitudinally adjacent XPP cells specifies lateral root founder cells (LRFCs) (Torres-Martínez et al., 2020). When two longitudinally adjacent LRFCs are specified, their nuclei migrate toward their common cell wall prior to asymmetric cell division (Goh et al., 2012). This asymmetric cell division produces daughter cells with two distinct identities. Genes regulated by the transcription factor LATERAL ORGAN BOUNDARIES-DOMAIN 16 (LBD16) are required for auxin-induced polar nuclear migration and subsequent asymmetric cell division (Goh et al., 2012). It is also known that F-actin is required for the polar nuclear migration of LRFCs (Vilches Barro et al., 2019). However, the mechanism of cell polarity establishment in LRFCs remains unknown.

To understand the role of auxin in LR initiation, high-resolution analysis of auxin dynamics is indispensable. Auxin dynamics during lateral root primordium (LRP) formation have long been studied using reporter systems in which reporter genes are expressed under the control of the auxin-responsive synthetic DR5 promoter (Dubrovsky et al., 2008; Moreno-Risueno et al., 2010). Since auxin reporters driven by auxin-inducible promoters depend on protein accumulation, there is an inherent time lag in the detection of auxin levels. Furthermore, the high stability of GFP can result in a dissociation of reporter signals from dynamic auxin levels, especially when auxin levels are declining. While destabilized luciferase can track decreasing auxin levels, its spatial resolution is insufficient to distinguish single cells or specific LRP stages (Moreno-Risueno et al., 2010, Toyokura et al., 2019, Gernier et al., 2025). The R2D2 auxin reporter, which depends on auxin-inducible degradation of the DII domain, offers high spatiotemporal resolution (Liao et al., 2015). However, the RPS5A promoter, which was used in the original R2D2 is weak in mature XPP cells and only becomes strong after LRP formation begins. Here, we developed a fast-maturing, XPP/early LRP-specific R2D2 auxin sensor (fx-R2D2) tailored for investigating LR initiation with high spatiotemporal resolution.

Inhibition of PIN-mediated auxin transport decreases auxin levels within roots, except for the root tip, and consequently inhibits LR initiation. This decrease is caused by a reduced supply of auxin from the regions where auxin is produced (Reed et al., 1998; Casimiro et al., 2001). This suggests that auxin maxima are not formed when basal auxin levels fall below a specific threshold. Because inhibition of PIN function alone decreases the basal auxin level below the threshold, it does not allow us to properly assess the role of PINs in the formation of auxin maxima and the LRP. To circumvent this problem, Benková et al. examined the effects of a PIN inhibitor or mutations in *PIN* genes in the presence of exogenous auxin supply (Benková et al., 2003). External auxin treatment synchronously activates LRP formation throughout the pericycle. Under such conditions, a PIN inhibitor or mutations in multiple *PIN* genes cause unorganized cell division in the pericycle along the root without LRP formation. This indicates the importance of local PIN function in LRP formation. However, it remains to be determined which specific stage of LRP formation is disrupted by PIN inhibition.

Here, using fx-R2D2 live imaging, we unequivocally demonstrate the spatiotemporal auxin dynamics during LR initiation. Furthermore, fx-R2D2 imaging, together with nuclear tracking upon external auxin treatment, reveals that PINs regulate initial auxin patterning and coordinated polar nuclear migration.

## Materials and Methods

### Plant Material and growth conditions

Wild type Columbia-0 (Col-0) seeds and original R2D2 reporter line (Liao et al., 2015) were obtained from the Arabidopsis Biological Resource Center (http://www.arabidopsis.org). All wild type transgenic lines made in this study were of the Col-0 background. The *pin3 pin7* mutant line (Benková et al., 2003), *PIN1 promoter:PIN1-GFP* line (Benková et al., 2003) and *PIN7 promoter:PIN7-GFP* line (Blilou et al., 2005) were kindly provided by J. Friml. *PIN3 promoter:PIN3-GFP* line (Ioio et al., 2008) was kindly provided by M. Terao-Morita. *DR5 promoter:NLS-Venus* line (Heisler et al., 2005) was kindly provided by E. Meyerowitz. *PFA2 promoter:HISTONE H2B-tdTomato* construct and Col-0 transgenic plants carrying this reporter were generated in a previous report (Zhang et al., 2021). For *pin3 pin7* mutant experiment, *PFA2 promoter:HISTONE H2B-tdTomato* construct was transformed into the *pin3 pin7* mutant plants using floral dip method (Clough et al., 1998).

The growth medium used in this study (1/5MS medium) contained 0.2x MS medium salt (Murashige and Skoog, 1962), 1% (w/v) sucrose, 0.05% (w/v) MES-KOH (pH 5.7), 0.01% (w/v) inositol, 0.002% (w/v) thiamin hydrochloride, 0.00002% (w/v) nicotinic acid, 0.00002% (w/v) pyridoxine hydrochloride. Solid media were solidified with 0.4% (w/v) Phytagel^TM^ (Sigma-Aldrich). For Phytagel^TM^-solidification, CaCl2·2H2O and MgSO4·7H2O were added to a final concentration of 1x MS in all media.

Seeds were sterilized with 70% ethanol on ice for 5 minutes and, in some experiments, subsequently treated with 1/6x diluted sodium hypochlorite solution (nacalai tesque, Kyoto, Japan) at 4 °C for 10 minutes before washing with cold sterilized water; and then kept at 4 °C for 1 to 10 nights before use. Sterilized seeds were grown on solid medium under continuous light at 23 °C.

### Time-lapse conditions

5-day-old seedlings were placed on the glass surface of a 20 x 45 mm glass-bottom chamber (Cat No 5202-001, IWAKI, Shizuoka, Japan) wetted with 200 µL of liquid medium containing 0.02 % (v/v) of dimethyl sulfoxide (DMSO) with or without 2 µM Indole-3-acetic acid (IAA) and 10 µM naphthylphthalamic acid (NPA). Subsequently, the root region was overlaid with the same medium solidified with 0.4% (w/v) Phytagel. IAA and NPA stock solutions were prepared in DMSO and added to the autoclaved medium, maintaining a constant final DMSO concentration of 0.02% (v/v) across all treatments.

Fluorescence microscopic z-stack images were taken every 10 minutes for 24 hours using an Axio Observer Z3.1 (ZEISS) fluorescence microscope and Axiocam 506 mono (ZEISS) CCD camera (for PFA2 reporters and original R2D2 reporter) or ORCA Flash4.0 (Hamamatsu Photonics) CMOS camera (for fx-R2D2 reporters) under dark conditions. Within the 10-minute period, at most 5.5 minutes were spent acquiring images, and during the remaining time the entire seedlings were exposed to the light from a halogen lamp to enable photosynthesis.

### Quantification of polar nuclear migration under auxin treatment

*PFA2 promoter:HISTONE H2B-mRuby3* construct and corresponding transgenic reporter line were generated in this study (Supplementary Material S1). Regions located approximately 1.2–2.9 mm from the root tip at the start of time-lapse imaging were defined as regions of interest (ROIs). ‘Reference positions’, ‘division positions’ and ‘directions of nuclear migration’ were determined by using the method explained in Results and Fig. 3.

### Quantification of auxin response

The *fx-R2D2* plasmid containing *PFA2 promoter:DII-NLSmScarlet-I* and *PFA2 promoter:mDII-NLSmNeonGreen* was generated in this study (Supplementary Material S2). The 3181 bp region immediately upstream of the translation start site of the *PFA2* gene was used as the *PFA2* promoter. *mScarlet-I* and *mNeonGreen* were codon-optimized for *Arabidopsis thaliana* (Qian et al., 2018). The construct was transformed into wild type plants. Fluorescence microscopic images were taken under the conditions described above. The same regions used in the nuclear migration analysis were selected as ROIs. Data processing and analysis were performed using Fiji ImageJ software. Time-lapse images were first processed with Median filter with a parameter of radius = 2 pixels. Background signals were subtracted from the obtained images by Subtract Background tool. DII/mDII ratio calculation was done with Calculator Plus plugin (Divide). To extract nuclear regions, mDII images were processed with a median filter followed by background subtraction as described above. Processed mDII images were then thresholded to generate binary masks corresponding to nuclear regions. DII/mDII ratio images were multiplied by the binary masks using the Calculator Plus plugin (Multiply) to extract nuclear signals. Maximum-intensity projection images were subsequently generated using the Z Project tool (Max Intensity).

### PIN localization analysis

Wild type plants carrying *PIN promoter:PIN-GFP* reporters were crossed with plants carrying *PFA2 promoter:HISTONE H2B-tdTomato* reporter, and F3 plants containing both reporters were used for observation. 5-day-old seedlings were fixed with a fixative solution (2.5% (w/v) paraformaldehyde, 1% (v/v) DMSO, 0.1% (v/v) Triton X-100, 0.2% (v/v) SCRI Renaissance Stain 2200 (SR2200) (Renaissance Chemicals, Selby, UK) (Musielak et al., 2015), 1x PBS buffer (pH 7.4)) for 30 minutes at room temperature, and washed with ClearSee (Kurihara et al., 2015) for clearing. The cleared samples were mounted with ClearSee and observed using an FV1000 (Olympus, Tokyo, Japan) confocal laser scanning microscope (CLSM). Z-stack images were stitched using the Pairwise Stitching plugin in Fiji ImageJ. The LRFC-containing regions closest to the root tip were observed in each sample.

## Results

### High spatiotemporal observation of auxin dynamics during LRP initiation using a new auxin reporter

To reveal auxin dynamics with high spatiotemporal resolution, we developed an XPP/early LRP-specific R2D2 auxin reporter using fast-maturing fluorescent proteins (fx-R2D2). The widely used auxin reporter R2D2 reflects auxin levels based on the ratio of an auxin-sensitive DII-nVenus signal to an insensitive mDII-ntdTomato signal (Liao et al., 2015). Here, we employed DII fused to nuclear-localized mScarlet-I (DII-NLSmScarlet-I) and mDII fused to nuclear-localized mNeonGreen (mDII-NLSmNeonGreen), expressed from transgenes driven by the XPP/early LRP specific *PFA2* promoter. Under normal growth conditions, the DII signal compared with the mDII signal was high from the root meristem to the beginning of the elongation zone, indicating relatively low auxin levels in XPP cells in this region. Auxin levels increased in a broad region on the convex side of the bending root. (1h 50 min in Fig. 1, Supplementary Video S1). Subsequently, the DII signal quickly decreased further to an undetectable level in a subset of XPP cells (2 h 50 min in Fig. 1, Supplementary video S1), followed by polar nuclear migration (3 h 40 min to 4 h 30 min in Fig. 1) and asymmetric cell division (6 h 20 min in Fig. 1). The high auxin level was maintained during LRP development (2 h 50 min – 11 h 20 min in Fig. 1, Supplementary Videos S1). When two LRPs began to form in close proximity, auxin levels rapidly decreased in the cells of one of the LRPs, and cell division subsequently ceased in these cells (cells containing nuclei 5′ and 6′ in Fig. 1, Supplementary Video S1). These findings indicate that changes in auxin levels quickly determine whether cells undergo subsequent division.

**Figure 1.**
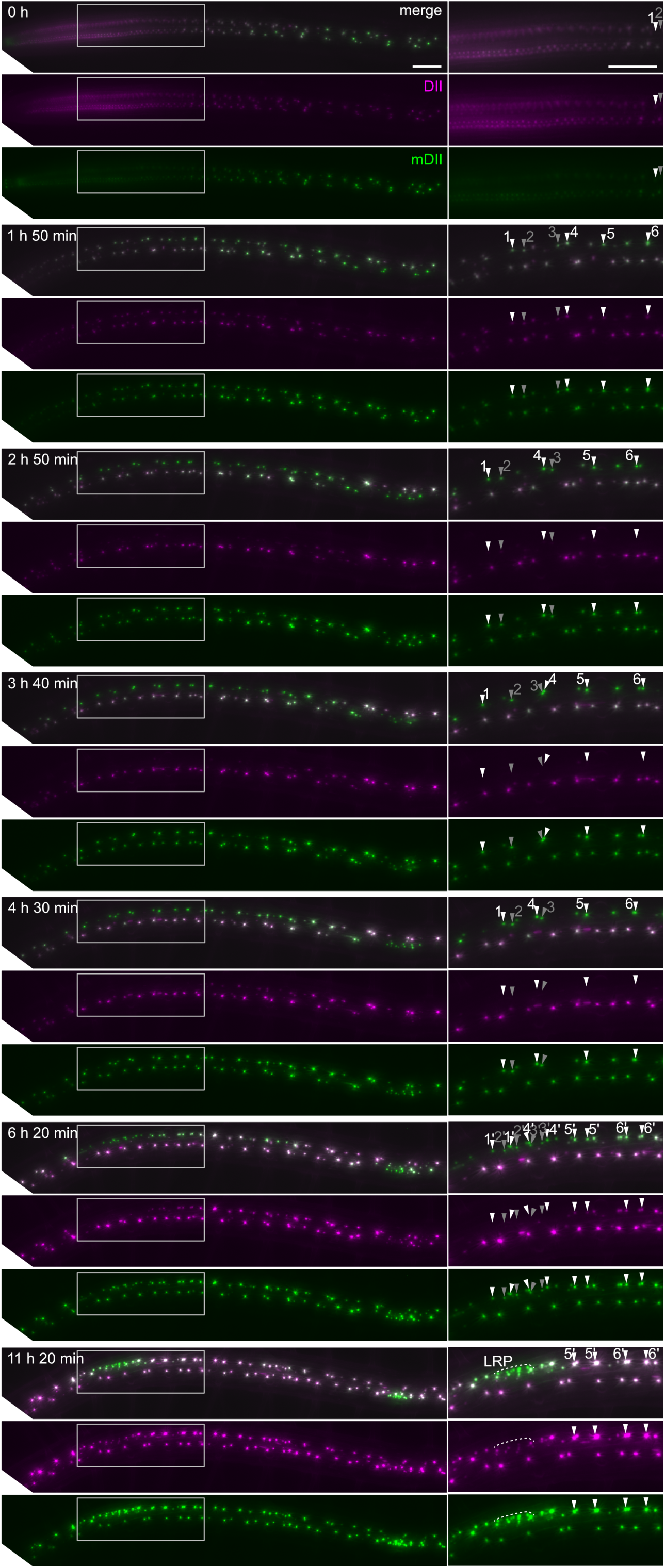
High-resolution live imaging of auxin dynamics during LR initiation and early LRP development. Fluorescence microscopic time-lapse images of a 5-day-old seedling expressing *fx-R2D2* overlaid with 1/5MS medium. The corresponding time-lapse movie is provided as Supplementary Video S1. For each time point, the images are arranged from top to bottom as follows: merged maximum-intensity projection images of DII-NLSmScarlet-l and mDII-NLSmNeonGreen, maximum-intensity projection images of DII-NLSmScarlet-l, maximum-intensity projection images of mDII-NLSmNeonGreen. Right panels show enlarged views of the regions outlined by the gray rectangles in the left panels. White and gray arrowheads indicate XPP nuclei. Labels T-6’ indicate the daughter nuclei derived from nuclei 1-6, respectively. The numbers shown in the upper right corners of the merged images in the left panels (h min) indicate the elapsed time from the start of the time-lapse observation, which was defined as 0 h in the top panel. Scale bars = 100 pm. Cells in a broad region of the XPP cell files on the convex side of the root bend exhibit relatively high auxin levels. Groups of cells in this region rapidly decrease their Dll signal (2 h 50 min) and subsequently undergo polar nuclear migration (from 3 h 40 min to 4 h 30 min) and asymmetric cell division (6 h 20 min), giving rise to stage-l LRPs. LRPs that maintain high auxin levels continue to grow (nuclei 1 – 4’), whereas those that decrease their auxin levels (6’ and 7’) cease cell division.

### PINs are required for auxin patterning under external auxin treatment

To investigate the role of PINs in initial auxin patterning in LRFCs, we tested auxin distribution during LR initiation and the early stages of LRP formation under external auxin treatment. We first examined the expression pattern of auxin reporter *DR5 promoter:N7-Venus* (Heisler et al., 2005) under an IAA 2µM treatment. The signal appeared to be present in all cells and had no specific pattern (Supplementary Figure S1). *DR5 promoter:N7-Venus* is an excellent reporter when auxin response is constant or increasing. However, due to the high stability of Venus, this reporter may not accurately reflect the actual auxin response when auxin response is decreasing or fluctuating. Therefore, we used fx-R2D2 to examine auxin status under externally auxin application. 2 µM IAA treatment quickly decreased the DII signal to undetectable levels (Fig. 2b). After four hours of IAA treatment, the signal recovered in the regions where LRPs did not form but remained low at the sites that later developed into LRPs (Fig. 2j, Supplementary Figure S2). This result indicates that auxin response is quickly and strongly induced in all XPP cells, and subsequently, it diminishes in XPP cells that do not undergo LRP formation, even under continuous auxin treatment. Importantly, NPA impaired the formation of this pattern (Fig. 2m-p, Supplementary Figure S2). This result suggests that PIN-driven polar auxin transport is required for auxin response patterning in the early stages of external auxin-induced LRP formation.

**Figure 2.**
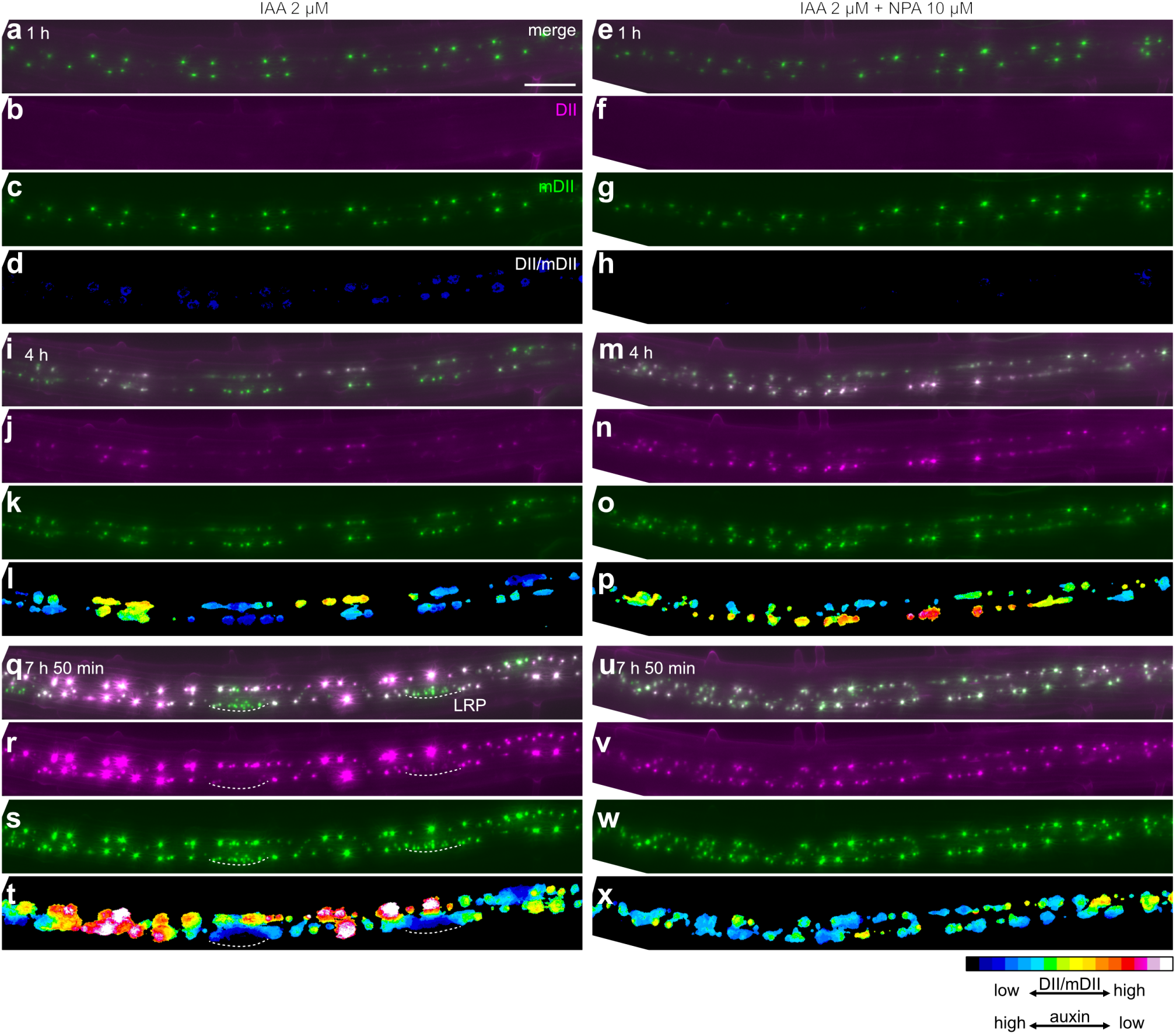
*f×-R2D2* signal patterns under 2 pM IAA treatment. Fluorescence microscopic time-lapse images of 5-day-old seedlings expressing fx-R2D2. The seedlings were treated with 2 pM IAA (left) or 2 pM IAA + 10 pM NPA (right). Five plants were used for each treatment, and a representative plant is shown for each. DIl/mDIl-ratio images for all plants are shown in Supplementary Figure S2. Merged maximum-intensity projection images of DII-NLSmScarlet-l and mDII-NLSmNeonGreen (a, e, i, m, q, u), individual channel maximum-intensity projection images of DII-NLSmScarlet-l (b, f, j, n, r, v), mDII-NLSmNeonGreen (c, g, k, o, s, w) and maximum-intensity projection images of DII/mDII ratio signals within nuclear regions (d, h, I, p, t, x)are shown. Images were acquired after 1h (a-h), 4h(i-p), 7h50min (q-x) of each treatment. Regions located approximately 1.2-2.4 mm from the root tip at the start of time-lapse imaging are shown. Scale bars = 100 pm. Regardless of the presence of NPA, 1 h of IAA treatment reduced the Dll signal to an undetectable level (b, f). After 7 h 50 min of IAA treatment, LRPs with a high auxin response had formed (q-t), whereas no distinct LRPs were observed after 7 h 50 min of IAA + NPA treatment (u-x).

### Auxin treatment synchronously induced polar nuclear migration of XPP cells followed by asymmetric cell division

Next, we tested whether polar nuclear migration is induced in XPP cells by 2 µM IAA treatment. We used Arabidopsis seedlings expressing *HISTONE H2B-tdTomato* reporter or *HISTONE H2B-mRuby3* reporter under the control of the XPP/early LRP-specific *PFA2* promoter to monitor the migration of XPP nuclei by time-lapse imaging. In our experimental conditions, all XPP nuclei divided within 5 hours of the treatment (Supplementary Figure S3). For quantification, we first determined the center positions of XPP nuclei at 1 hour after the start of IAA treatment (hereafter, we refer to such positions as ‘reference positions’). The distances between all neighboring reference positions within each cell file (black double-headed arrows in Fig. 3a) showed a symmetric, bell-shaped distribution around a median of 137.4 µm (Fig. 3b, left), indicating that nuclei are not randomly positioned, but tend to be evenly spaced.

**Figure 3.**
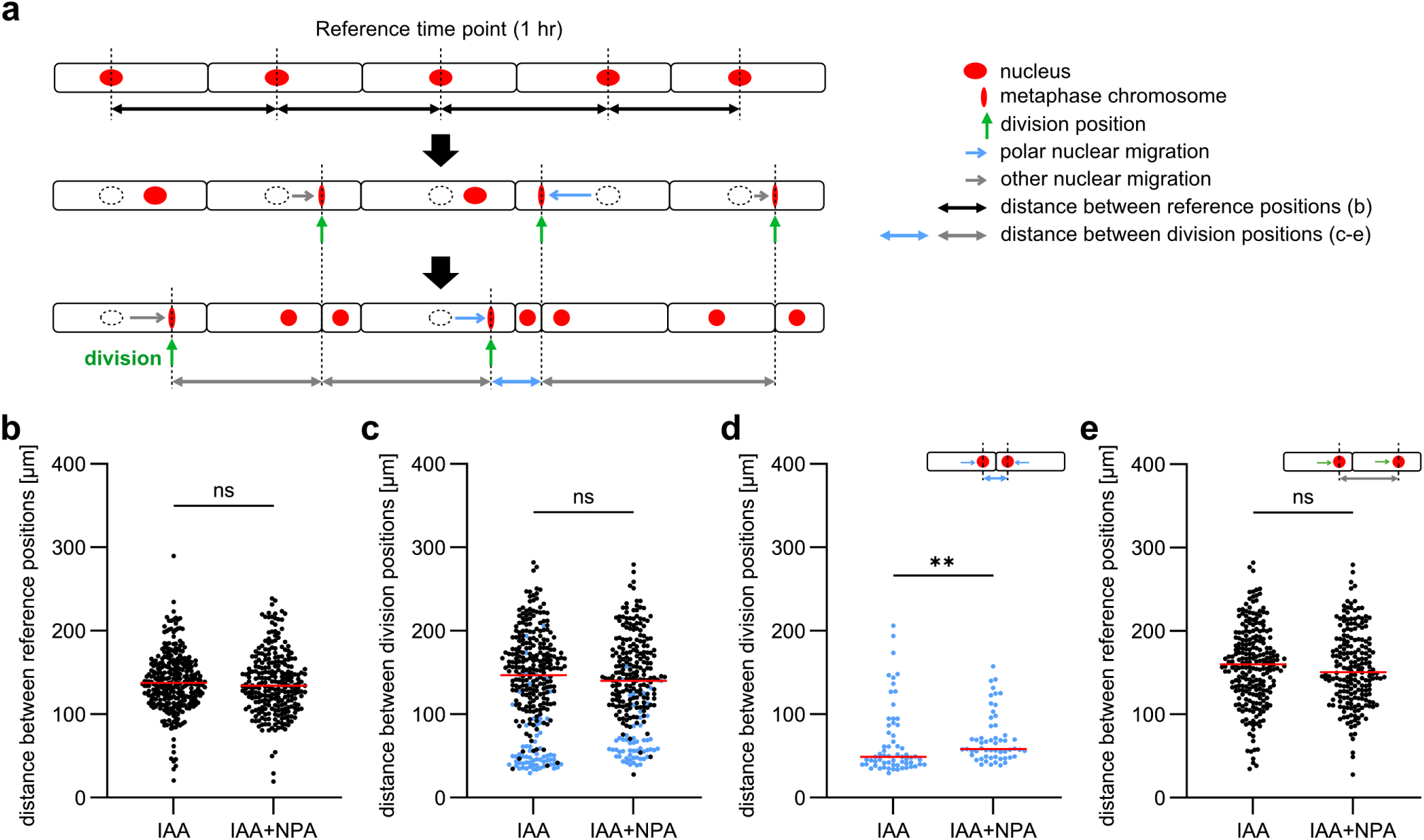
The PIN inhibitor NPA impairs polar nuclear migration of XPP cells. **a.** A schematic for analysis of auxin-induced nuclear migration in XPP. Each rectangle indicates an individual XPP cell. The centers of XPP nuclei at 1 hour after the start of 2 pM IAA treatment were set as the ‘reference positions’. The division positions were defined as the positions of the metaphase plate (green arrows). The distances between division positions of all longitudinally adjacent XPP cells were determined (blue and gray double-sided arrows). Longitudinally adjacent nuclei pairs that migrated toward each other and subsequently underwent synchronous nuclear division within 40 min were classified as exhibiting polar migration (blue arrows), **b.** Distances between reference positions of all longitudinally adjacent XPP cells, n = 305 (IAA), 258 (IAA + NPA), c. Distances between division positions of all longitudinally adjacent XPP cells. Blue dots represent distances between division positions of nuclei pairs with polar migration, n = 305 (IAA), 258 (IAA + NPA), **d, e,** Blue and black dots from c are shown separately, n = 65 (IAA), 56 (IAA + NPA) (d), n = 240 (IAA), 202 (IAA + NPA) (e). Mann-Whitney test (**0.001 ≤ P < 0.01, *0.01 ≤ P < 0.05. ns, not significant (P ≥ 0.05)) was used to evaluate statistical significance (b-e).

Then, we marked the positions of the metaphase plates during the first round of cell divisions after IAA treatment (hereafter, we refer to such positions as ‘division positions’). Then, we measured the distances between division positions of longitudinally adjacent XPP cells in each cell file (blue or gray double-headed arrows in Fig. 3a). The distribution of the distances between the division positions showed a distinct peak around 40 µm (Fig. 3c, left).

We then determined the directions of nuclear migration by comparing the ‘reference positions’ and the ‘division positions’ of each nucleus (green or blue arrows in Fig. 3a). Based on these directions, we identified pairs of XPP nuclei that moved toward each other before the first cell division following IAA treatment (Supplementary Video S2). We defined ‘nuclei with polar migration’ based on the following criteria: 1. the nuclei approached each other before the first cell division, 2. they completed their division within a 40 min-time difference. The peak around 40 µm in Fig. 3c corresponded to XPP nuclear pairs with polar migration, most of which gave rise to future LRPs (Supplementary Figure S4).

### The PIN inhibitor impaired polar nuclear migration in XPP cells

To investigate the role of polar auxin transport in polar nuclear migration, we tested the effects of NPA, a PIN inhibitor (Ung et al., 2022; Yang et al., 2022). Under our experimental conditions, NPA did not affect the timing of auxin-induced XPP cell division (Supplementary Figure S3). However, NPA treatment partially but significantly increased the distances between division positions of adjacent nuclei undergoing polar migration (Fig. 3c, d). This result suggests that PIN-driven polar auxin transport is important for polar nuclear migration in LRFCs.

### PIN localization in LRFCs and their adjacent XPP cells

To understand the role of PINs in the polar nuclear migration, we examined the localization of PIN proteins in LRFCs and their longitudinally adjacent XPP cells. Among the six plasma membrane-localized PIN transporters, *PIN1*, *PIN3*, *PIN4* and *PIN7* are highly expressed in the XPPs, and *PIN1* and *PIN7* are induced by exogenous auxin treatment (Supplementary Figure S5, De Smet et al., 2008). PIN localization in LRPs and LRFC-overlying endodermal cells have been reported (Benková et al., 2003; Marhavý et al., 2013; Omelyanchuk et al., 2016). However, to our knowledge, their localization in LRFCs and their adjacent XPP cells have not been reported in detail. This was likely because it was difficult to identify LRFCs without an LRFC marker. Here, we used an XPP nuclear marker and *PIN promoter:PIN-GFP* reporters to examine the localizations of PINs in LRFCs and their longitudinally adjacent XPP cells. Pairs of longitudinally adjacent XPP cells with an inter-nuclear distance of ≤ 70 µm, measured between the centers of the nuclei, were defined as LRFCs. PIN1, PIN3 and PIN7 were highly localized to the plasma membrane of LRFCs on the side facing their common cell wall. In addition, PIN7 was also localized to the plasma membrane of longitudinally adjacent XPP cells on the side facing the LRFCs (Fig. 4b-j, Supplementary Figure S6-10, Table 1). PIN3 was occasionally localized to the plasma membrane of XPP cells basally adjacent to the basal LRFC (Fig. 4e-g, Supplementary Figure S7, Table 1). These localization patterns suggest that PIN1, PIN3 and PIN7 may mediate auxin efflux toward the common cell wall between adjacent LRFCs, while PIN 7 and PIN3 may also contribute to auxin accumulation in LRFCs from adjacent XPP cells.

**Figure 4.**
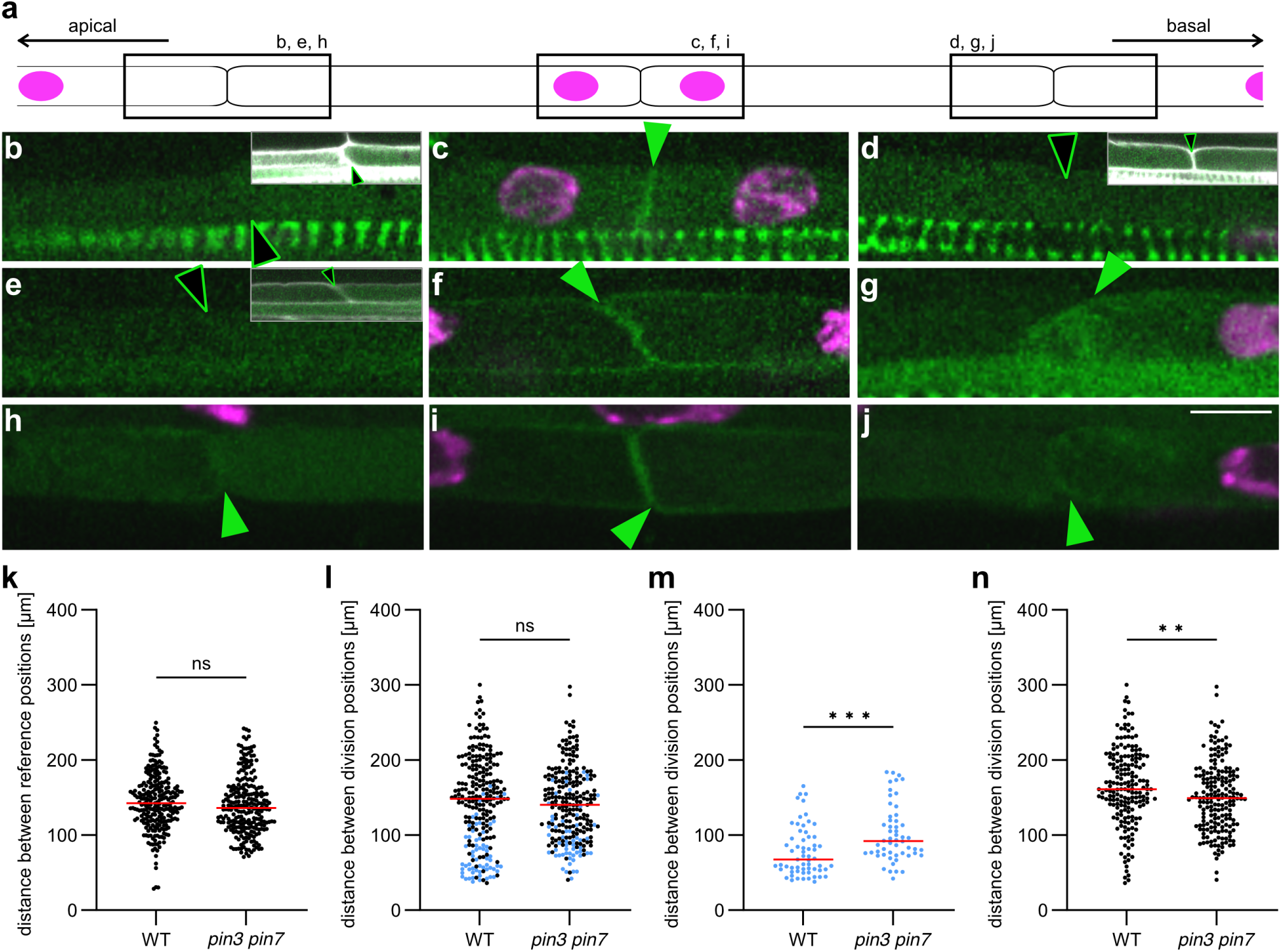
PIN3 and PIN7 contribute to the polar nuclear migration in XPP cells. **a.** An overview of the observed region in this experiment. The border between two LRFCs (center) and between a LRFC and its longitudinally adjacent XPP cell (both sides) were observed, **b-j.** Enlarged images of the regions shown with squares in a. The localizations of PIN1-GFP (b-d), PIN3-GFP (e-g) or PIN7-GFP (h-j) in wild type 5-day-old seedlings were observed with CLSM after fixation. Magenta, green and white represent signals from XPP nuclear marker *PFA2 promoter: HI STONE H2B-tdTomato, PIN promotenPIN-GFP* and cell wall stained with SR2200 respectively. Only magenta and green signals were shown in the large images. Small images at the upper right of the pictures show cell wall signal of the same regions of the large images. Green arrowheads indicate the positions of anticlinal cell interfaces with GFP signal, while the open arrowheads indicate the anticlinal cell interfaces lacking GFP signal. Scale bar = 10 pm. A summary of the PIN localization analysis is provided in Table 1. **k-n.** 5-day-old wild type (WT) and the *pin3 pin7* mutant seedlings expressing *PFA2 promoterHISTONE H2B-tdTomato* were treated with 2 pM lAAand 500 nM NPA. **k.** Distances between reference positions of all longitudinally adjacent XPP cells, n = 250 (WT), 248 *(pin3 pin7).* **I.** Distances between division positions of all longitudinally adjacent XPP cells. Blue dots represent distances between division positions of nuclei pairs with polar migration, n = 250 (WT), 248 *(pin 3 pin7).* **m, n.** Blue and black dots from I are shown separately, n = 61 (WT), 54 *(pin3 pin7)* (m), n = 189 (WT), 194 *(pin3 pin7)* (n). Mann-Whitney test (***P < 0.001, **0.001 ≤ P < 0.01, *0.01 ≤ P < 0.05. ns, not significant (P ≥ 0.05)) was used to evaluate statistical significance (k-n).

**Table 1.** Localization of PIN1, PIN3 and PIN7 at the boundary of LRFCs (the apical FC and the basal FC) and at the boundary of an apical or basal FC and the adjacent XPP cell.

| boundary | apical XPP / apical FC | apical FC / basal FC | basal FC / basal XPP |
| --- | --- | --- | --- |
| PIN1 (n = 9) |  |  |  |
| GFP present at the boundary | 2 | 7 | 1 |
| GFP assigned to apical-side cell | 0 | 3 | 0 |
| GFP assigned to basal-side cell | 2 | 6 | 0 |
| PIN3 (n = 8) |  |  |  |
| GFP present at the boundary | 1 | 5 | 2 |
| GFP assigned to apical-side cell | 0 | 2 | 0 |
| GFP assigned to basal-side cell | 1 | 3 | 2 |
| PIN7 (n = 9) |  |  |  |
| GFP present at the boundary | 6 | 9 | 7 |
| GFP assigned to apical-side cell | 6 | 7 | 0 |
| GFP assigned to basal-side cell | 0 | 9 | 7 |

If the GFP signal at the cell boundary was continuous with the longitudinal side of a cell, it was assigned to that cell. The signal was assigned to both cells when the signal was continuous with the longitudinal side of both cells.

### PIN3 and PIN7 contribute to the polar nuclear migration in XPP cells

Since PIN3 and PIN7 were localized at the plasma membrane of XPP cells adjacent to LRFCs, we next examined whether the polar nuclear migrations of LRFCs were impaired in the pin3 pin7 double mutants. Under IAA treatment alone, polar nuclear migration in the pin3 pin7 mutants showed no significant difference compared to the wild type (Supplementary Figure S9). However, polar nuclear migration was significantly impaired in the pin3 pin7 double mutants compared to the wild type when co-treated with IAA and a low concentration of NPA (Fig. 4k-n). This result suggests that polar auxin transport mediated by PIN3 and PIN7 contributes to the polar nuclear migration in LRFCs.

## Discussion

In the earliest steps of LRP formation in *Arabidopsis thaliana*, auxin is spontaneously accumulated in several XPP cells, which defines these cells as LRFCs. The nuclei of two adjacent LRFCs migrate toward the common cell wall. The regulatory mechanisms for the auxin accumulation and the polar nuclear migration in LRFCs are not well understood. To enable high spatiotemporal resolution live imaging of auxin dynamics during LR initiation and early LRP development, we modified the R2D2 auxin reporter. In the original version, the *RPS5A* promoter was used to drive the expression of the reporter components (Liao et al., 2015). The *RPS5A* promoter activity is weak in the pericycle cells but strongly increases during LRP formation (Supplementary Figure S10). The fluorescent protein tdTomato, which was fused to mDII in the original report, has a relatively long maturation time (maturation half time of approximately 90 minutes *in vivo* at 32**°**C (Guerra et al., 2022)). In contrast, Venus, which is fused to DII in the original R2D2 reporter (Liao et al., 2015), has a relatively short maturation time (Nagai et al., 2002). This difference could potentially distort the estimation of auxin levels when DII and mDII expression is rapidly elevated. Here, we used the XPP/early LRP-specific *PFA2* promoter and fast-maturing fluorescent proteins, mScarlet-I (Bindels et al., 2016) and mNeonGreen (Shaner et al., 2013), which were fused to DII and mDII, respectively. We observed a rapid increase in auxin levels in a group of XPP cells, followed by polar nuclear migration in LRFCs (Fig. 1, Supplementary Video S1). Auxin levels remained high throughout LRP formation. In some closely spaced LRPs, we observed a rapid decrease in auxin levels in one LRP, after which cell division ceased (Fig. 1, Supplementary Video S1). These observations are consistent with previous findings obtained using DR5:luciferase (Toyokura et al., 2019, Gernier et al., 2025). However, the low spatial resolution of the DR5:luciferase reporter prevented the association of the luciferase signals with specific developmental stages of LRPs (Moreno-Risueno et al., 2010). In contrast, DR5:N7-Venus cannot accurately capture rapid decreases in auxin levels because of the high stability of Venus. Our study is the first to reveal cellular auxin dynamics during LR founder cell establishment, polar nuclear migration, subsequent development, and the developmental cessation of LRPs with high spatiotemporal resolution (Fig.1, Supplementary Video S1). 2 µM IAA treatment induced rapid degradation of DII in all XPP cells to undetectable levels (Fig. 2b). Notably, the signal recovered in the regions that subsequently did not develop into LRPs even in the presence of external IAA (Fig. 2j, Supplementary Figure S2) and this pattern formation was inhibited by NPA (Fig. 2m-p, Supplementary Figure S2). This result highlights the importance of regulation of auxin dynamics by PIN proteins in regions where LRPs do not develop.

By tracking all XPP nuclei under IAA treatment, we showed that external auxin induces synchronous polar nuclear migration followed by asymmetric cell division, and the auxin efflux carriers PIN3 and PIN7 are required for this polar nuclear migration. Inhibition of the PIN function impaired both auxin patterns and polar nuclear migration (Fig. 2, 3, 4, Supplementary Figure S2), suggesting that PIN-regulated initial auxin pattern is required for polar nuclear migration. The partial effect of NPA on inhibiting polar nuclear migration suggests that additional systems may also regulate this process. In Arabidopsis, auxin efflux is mediated by two types of auxin efflux carriers: PINs and ABCB/PGPs. NPA strongly inhibits PIN7 and partially inhibits PGP19 (Petrášek et al., 2006). It is possible that PGP auxin efflux carriers and/or AUX/LAX auxin influx carriers (Hammes and Pedersen, 2024) function together with PINs.

The PIN3-mediated auxin supply from the endodermis to the LRFCs has been suggested (Marhavý et al., 2013). Another study showed that PIN1 is re-localized to the upper side of the plasma membrane of the protoxylem cells in gravistimulation induced LR initiation, suggesting a protoxylem-mediated auxin supply (Ditengou et al., 2008). Here we showed that PIN7 accumulated at the plasma membrane of XPP cells adjacent to the LRFCs, and PIN3 was accumulated at the plasma membrane of some XPP cells basally adjacent to the founder cell (Fig. 4, Supplementary Figures S6-10, Table 1). These localization patterns suggest an auxin flow from adjacent XPP cells toward LRFCs, thereby generating the auxin accumulation at LRFCs. Furthermore, they were highly accumulated at the plasma membrane of LRFCs on the side facing their common cell wall (Fig. 4, Supplementary Figures S6-10, Table 1), which suggest auxin transport toward the cell wall shared between two LRFCs. A hypothetical accumulation of extracellular auxin between two LRFCs may guide polar nuclear migration through transmembrane kinases (TMKs), which function as extracellular auxin receptors (Huang et al., 2019), but this possibility remains to be addressed in future studies.

How auxin gradients are translated into polar nuclear migration remains an important question for future studies. One possibility is that the auxin gradient itself acts as a direct spatial cue to instruct cell polarity. Alternatively, auxin may trigger non-cell-autonomous signaling molecules that coordinate polarity between cells. Indeed, it is proposed that, in the shoot apical meristem, downstream targets of AUXIN RESPONSE FACTOR 5 / MONOPTEROS (MP) non-cell-autonomously direct PIN1 polarity toward MP-expressing cells (Bhatia et al., 2016). Given that high auxin levels are established in the pair of LRFCs and their nuclei migrate toward each other, it is plausible that auxin-inducible intercellular signaling molecules produced in these founder cells mutually attract their nuclei, thereby establishing cell polarity in a coordinated manner.

## Acknowledgments

We thank J. Friml (Institute of Science and Technology Austria) for providing the *pin3 pin7* double mutant seeds, *PIN1 promoter:PIN1-GFP* reporter line and *PIN7 promoter:PIN7-GFP* reporter line, M. Morita (National Institute of Basic Biology) for providing *PIN3 promoter:PIN3-GFP* reporter line and E. Meyerowitz (California Institute of Technology) for providing *DR5 promoter:N7-Venus* reporter. We also thank Y. Kondo (The University of Osaka) for valuable discussions and for providing a supportive research environment. Plasmid containing *HISTONE H2B* fused to codon-optimized *mRuby3* was generated by K. Kinoshita-Tsujimura.

## Author Contributions

T.K. and S.K. designed the research and wrote the paper. S.K. performed all the experiments and analysis.

## Abbreviations

CLSM: confocal laser scanning microscope
IAA: indole-3-acetic acid
LR: lateral root
LRFC: lateral root founder cell
LRP: lateral root primordium
NPA: naphthylphthalamic acid
XPP: xylem-pole pericycle

## Legends to supplementary Videos

**Video S1. *fx-R2D2* patterns during LR initiation and early LRP development under normal conditions**

Time-lapse movie of merged maximum-intensity projection images of DII-NLSmScarlet-I and mDII-NLSmNeonGreen shown in Figure 1. The numbers shown in the upper left corner (hh:mm) indicate the elapsed time from the start of the time-lapse observation, with the first frame defined as 00:00. Scale bar = 100 µm.

**Video S2. Polar Nuclear Migration under 2 µM of IAA treatment**

Fluorescence microscopic time-lapse images of 5-day-old seedlings carrying *PFA2 promoter:HISTONE H2B-mRuby3* (red) under 2 µM IAA treatment. numbers at the upper right indicate the duration of IAA treatment (hh:mm). arrowheads indicate XPP nuclei that showed polar nuclear migration. Scale bar = 50 µm.

## Description of supplementary Data

**Supplementary data S1. The plasmid sequence of *PFA2 promoter:HISTONE H2B-mRuby3***

Full-length plasmid sequence of *PFA2 promoter:HISTONE H2B-mRuby3* used in this study.

**Supplementary data S2. The plasmid sequence of *fx-R2D2***

Full-length plasmid sequence of *fx-R2D2* used in this study.

**Figure S1.**
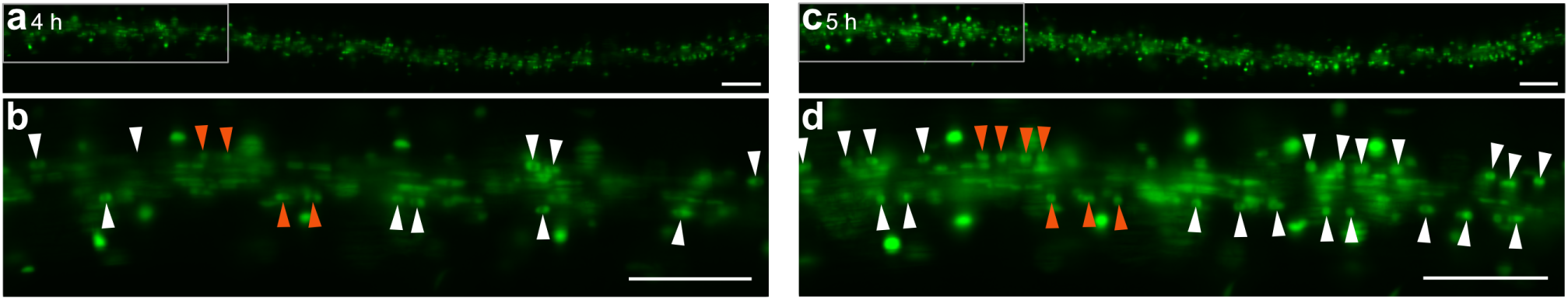
IAA induces DR5 signal without specific pattern. Fluorescence microscopic time-lapse images of 5-day-old seedlings carrying *DR5 promoterNLS-Venus* treated with 2 pM of IAA for 4 hours (a, b) or for 5 hours (c, d). b and d are enlarged images of the left side region of a and c indicated with gray rectangle, respectively. Arrowheads indicate pericycle nuclei. Orange arrowheads indicate nuclei of the pericycle that underwent asymmetric cell division under IAA treatment. Scale bar = 100 pm.

**Figure S2.**
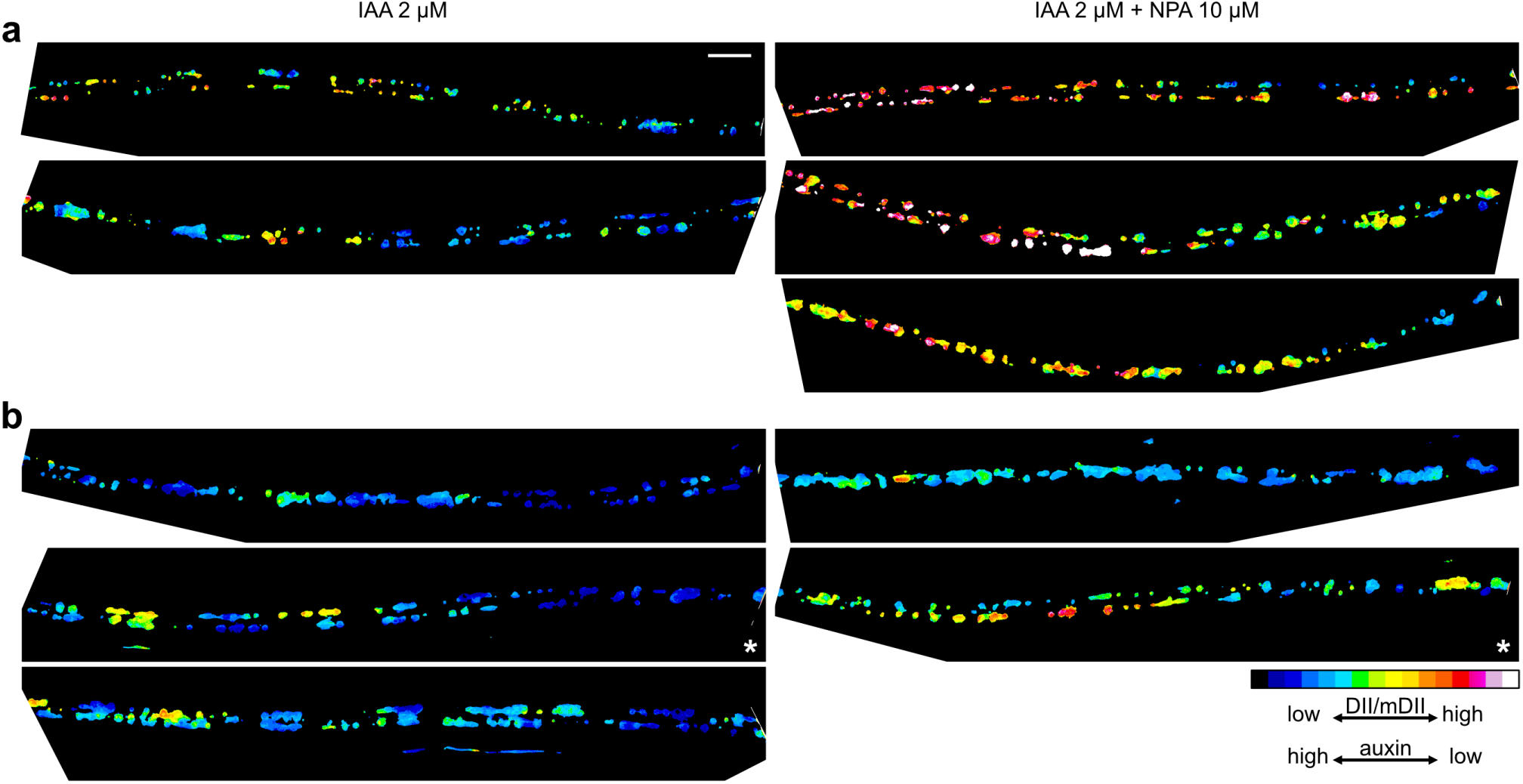
*f×-R2D2* signal patterns under 2 pM of IAA and 10 pM of NPA treatment. All seedlings observed in the experiment shown in Figure 2. Maximum-intensity projection images of DII/mDII ratio signals within nuclear regions at 4 hours of IAA treatment (left) or IAA + NPA treatment (right) are shown. Panels a and b represent two independent experiments performed on different days. For visualization purposes, the scaling coefficient (k1) was adjusted separately for each experiment. Regions located approximately 1.2-2.9 mm from the root tip at the start of time-lapse imaging were shown. Seedlings marked with an asterisk (*) in the lower right corner are the same individuals as those shown in Figure 2. Scale bar = 100 pm.

**Figure S3.**
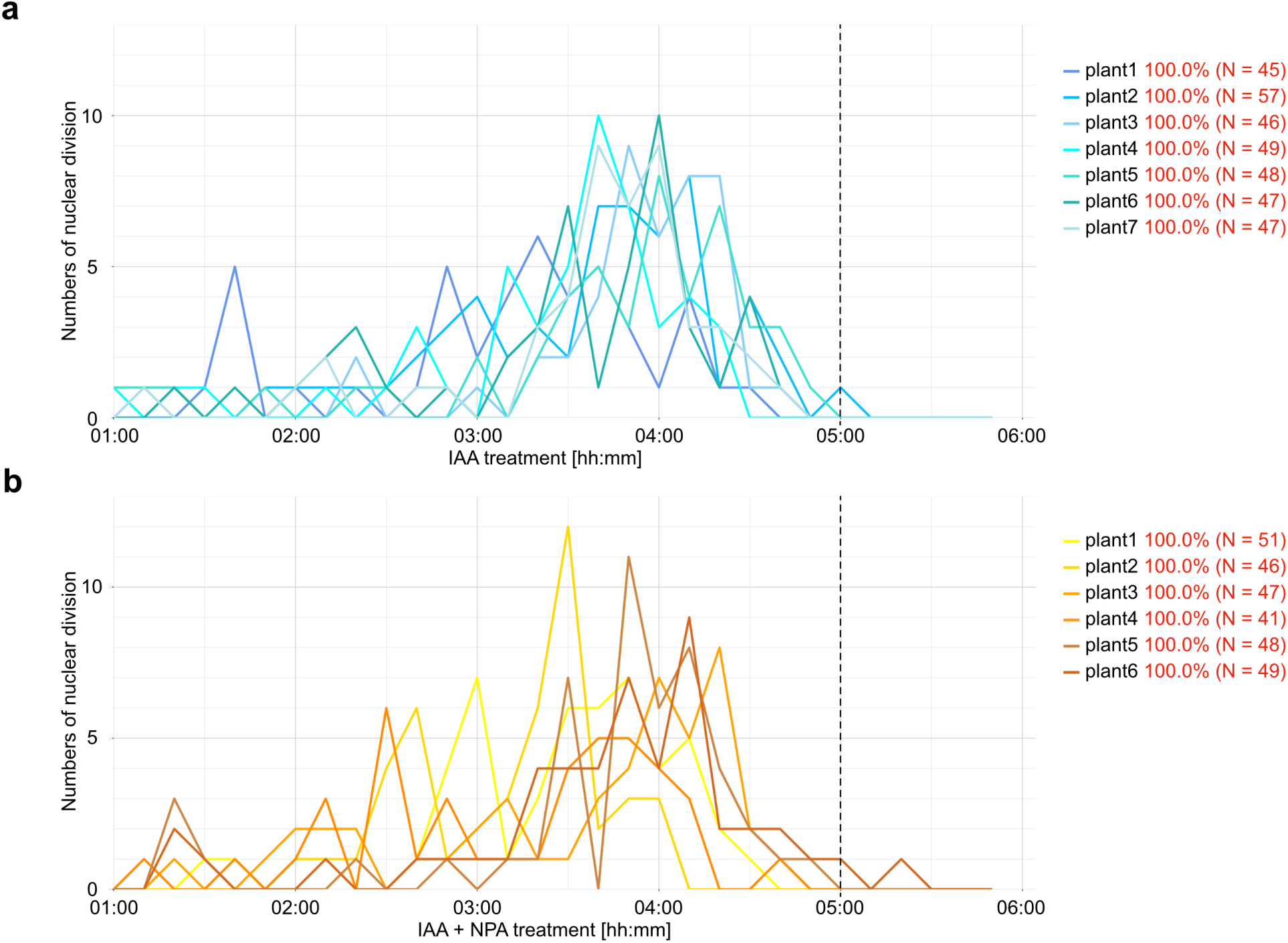
The timing of XPP cell division after external IAA treatment. The X axis shows the duration of 2 pM IAA (a) or 2 pM IAA + 10 pM NPA (b) treatment. The Y axis shows the numbers of nuclei that showed metaphase chromosome segregation at the indicated time points. Each line indicates independent plants used in Figure 3. Red numbers on the right side of each label indicate the percentage of nuclei that divided during the observation in each individual. All observed nuclei underwent their first cell division within 5 h 20 min. Mitotic cells at 1 h after IAA treatment were excluded from the analysis shown in Figure 3.

**Figure S4.**
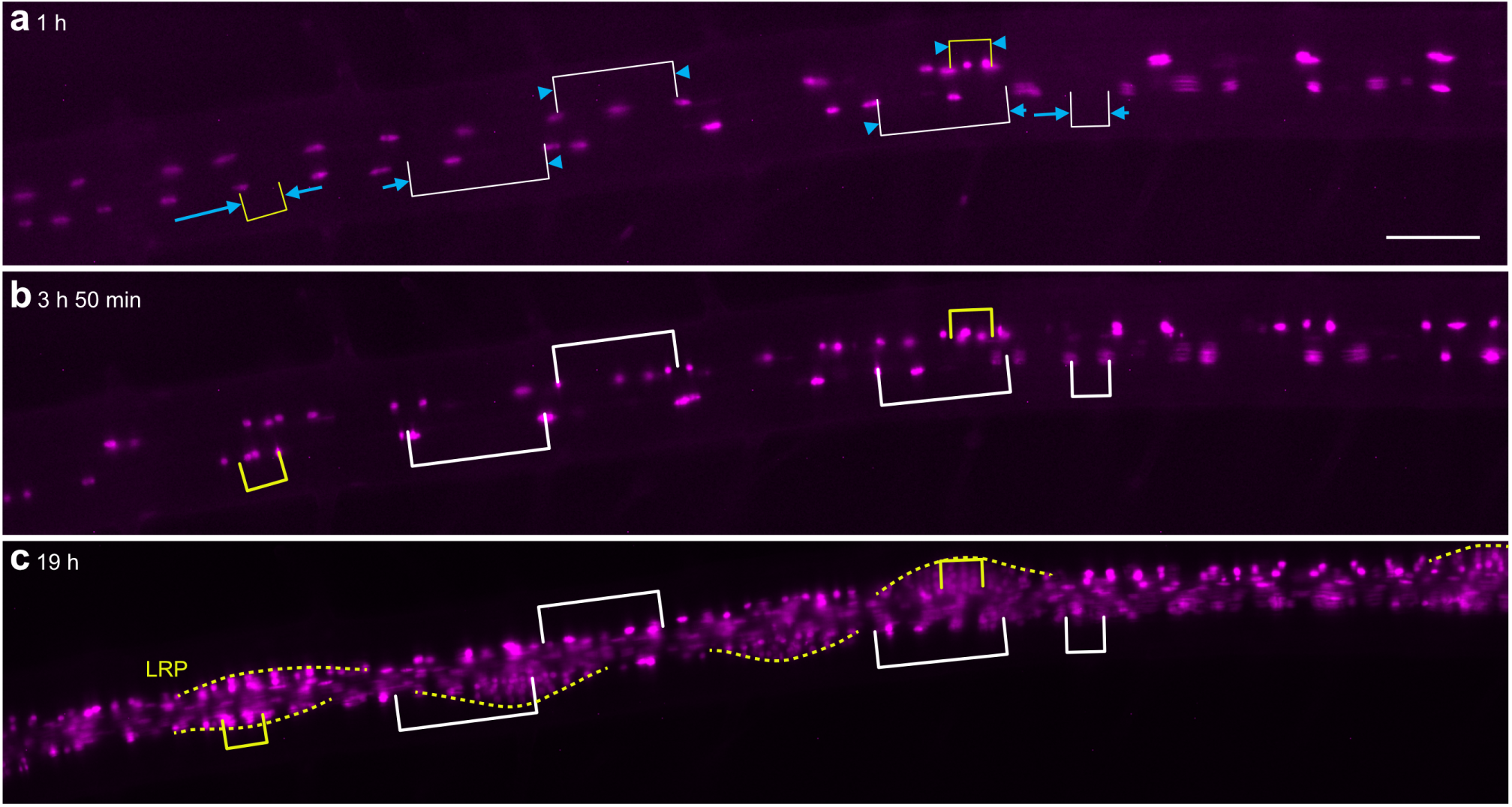
XPP-cell pairs whose nuclei migrated toward each other until the distance between them reached around 40 pm tend to give rise to LRPs. Fluorescence microscopic time-lapse images of 5-day-old seedlings carrying *PFA2 promoter: HI STONE H2B-mRuby3* (magenta) under 2 pM IAA treatment. Numbers at the upper left corners of each panel indicate the duration of IAA treatment (hh:mm). Blue arrows in panel a indicate the directions of each XPP nuclear migration before the first cell division after IAA treatment. White and yellow brackets indicate the distances between longitudinally adjacent division positions of the nuclei pairs that showed polar migration. Yellow brackets indicate the nuclei pairs of XPP cells that give rise to LRPs in c (yellow dashed line). Scale bar = 100 pm.

**Figure S5.**
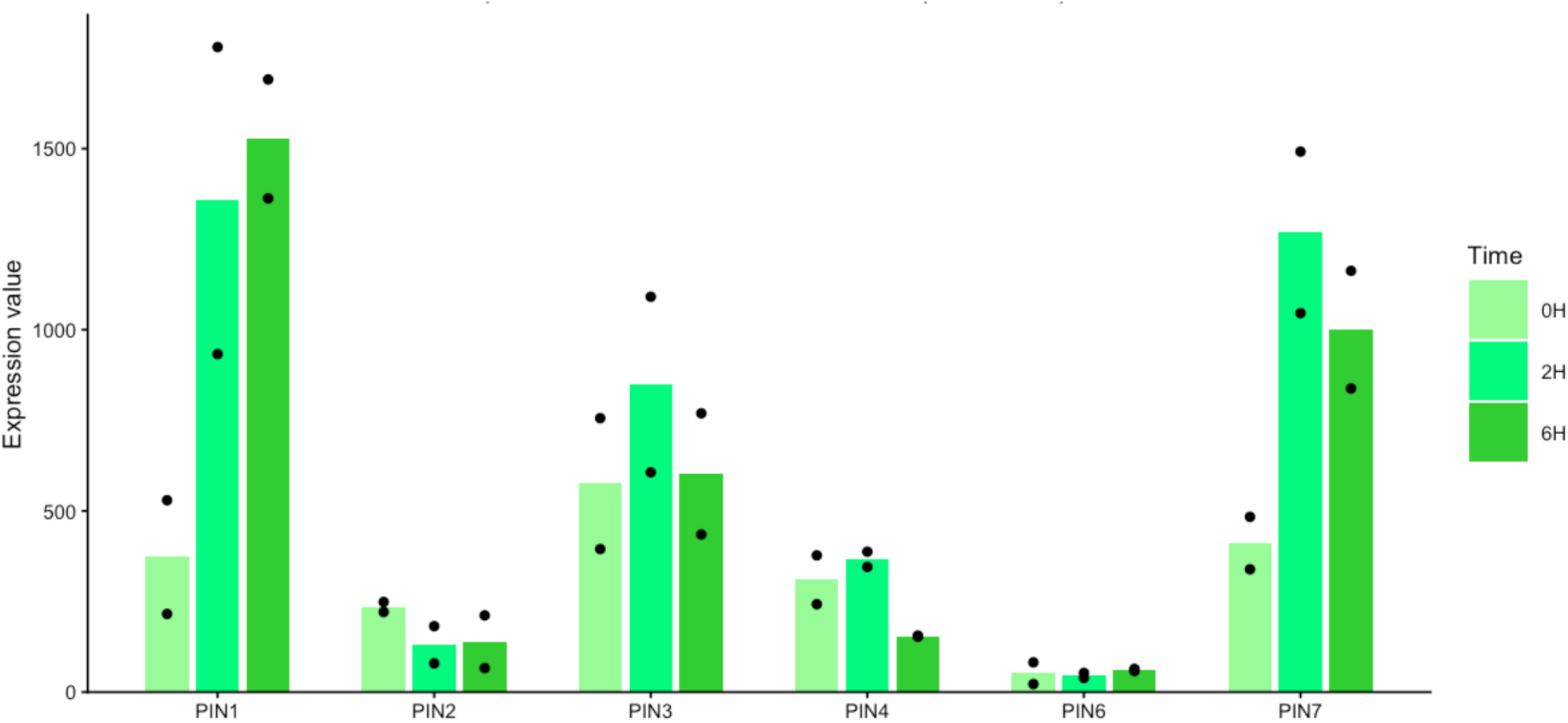
The expressions of *PIN* genes in XPP cells. Expressions of *PINs* in XPP cells in published microarray data sets (GSE6349, De Smet et al., 2008). Dots indicate expression values from two biological replicates. Each bar shows the average of two replicates.

**Figure S6.**
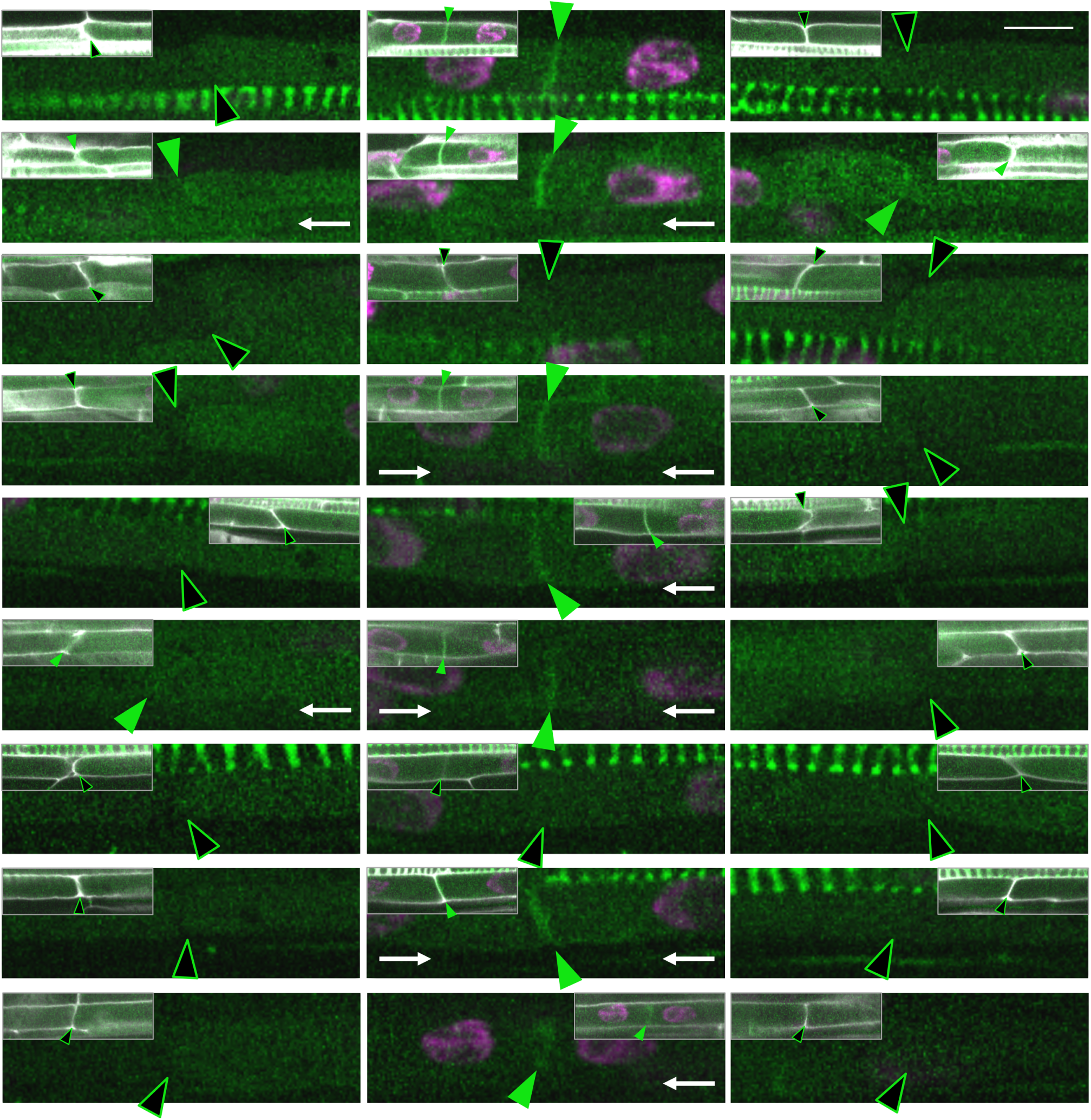
The subcellular localization of PIN1-GFP in the LRFCs and their longitudinally adjacent XPP cells. All confocal images used in Table 1 are shown. Magenta, green and white represent signals from XPP nuclear marker *PFA2 promotenHISTONE H2B-tdTomato, PIN1 promoter:PIN1-GFP* and cell wall stained with SR2200 respectively. Symbols shown below each image correspond to the cells on the respective sides of the image; the symbols on the left and right indicate data for the left and right cells, respectively. White arrows indicate anticlinal plasma membranes in which the GFP signal extended to the longitudinal side of the analyzed cell. Scale bar = 10 pm.

**Figure S7.**
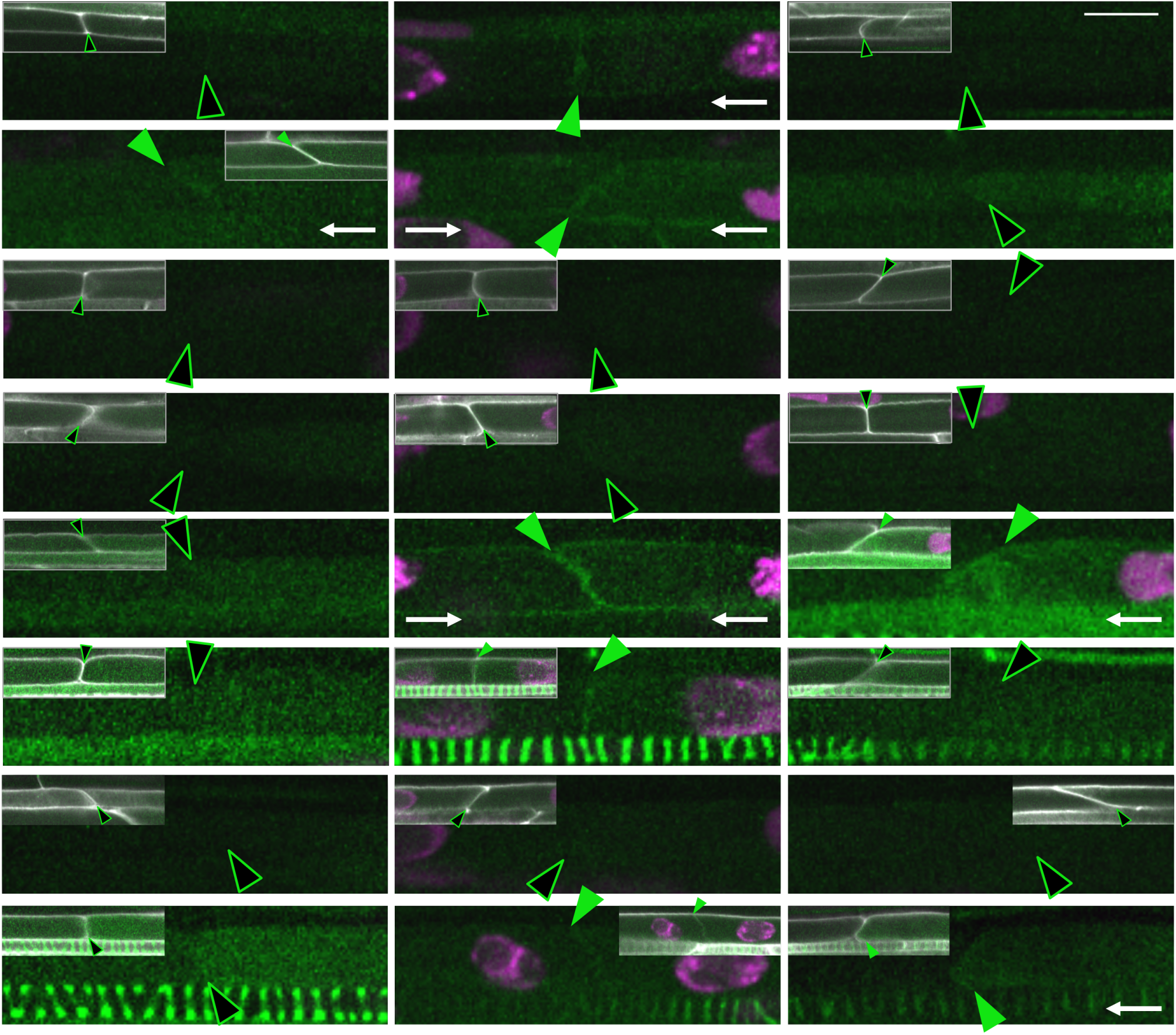
The subcellular localization of PIN3-GFP in the LRFCs and their longitudinally adjacent XPP cells. All confocal images used in Table 1 are shown. Magenta, green and white represent signals from XPP nuclear marker *PFA2 promoterHISTONE H2B-tdTomato, PIN3 promoter:PIN3-GFP* and cell wall stained with SR2200 respectively. Symbols shown below each image correspond to the cells on the respective sides of the image; the symbols on the left and right indicate data for the left and right cells, respectively. White arrows indicate anticlinal plasma membranes in which the GFP signal extended to the longitudinal side of the analyzed cells. Scale bar = 10 pm.

**Figure S8.**
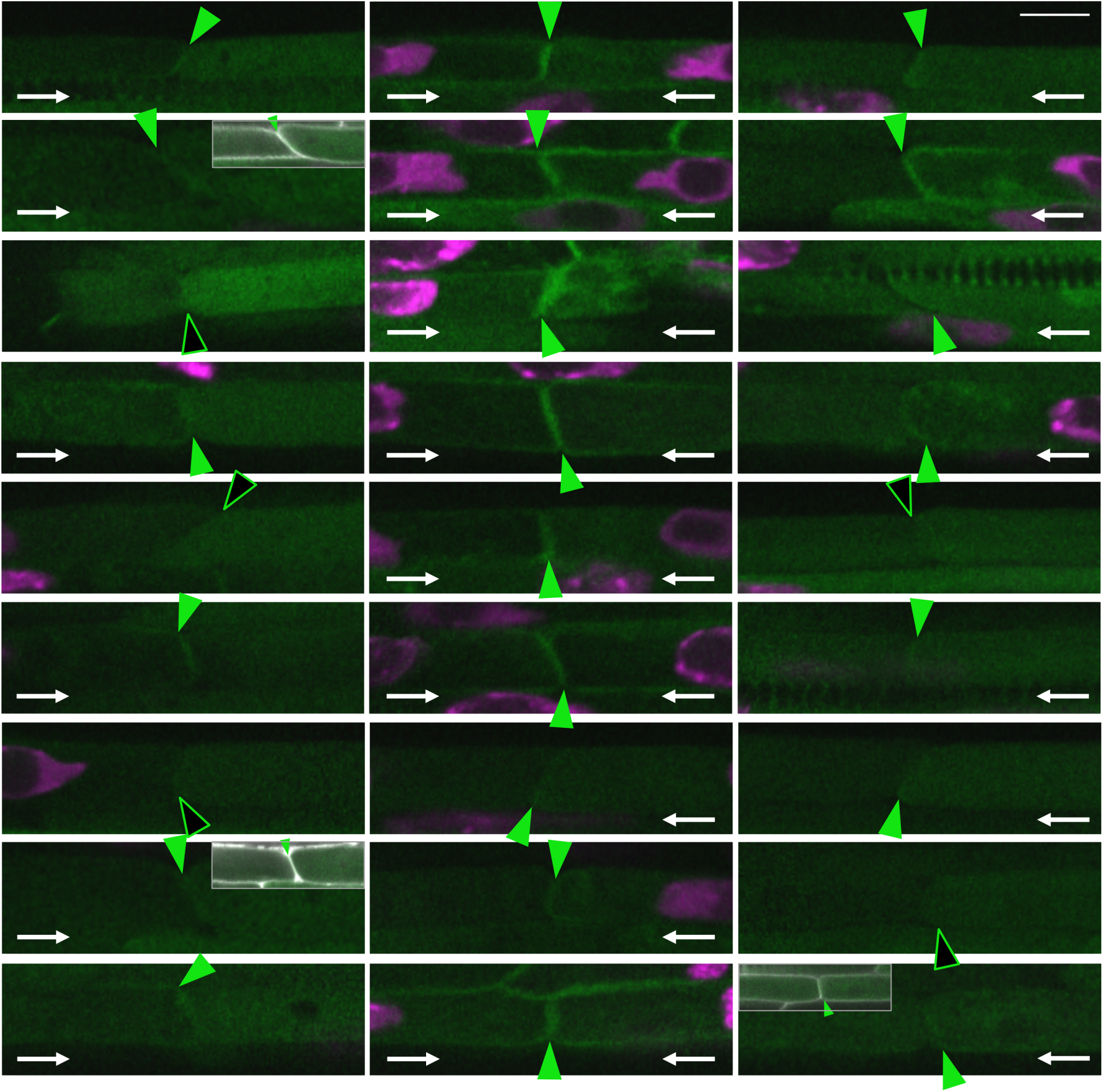
The subcellular localization of PIN7-GFP in the LRFCs and their longitudinally adjacent XPP cells. All confocal images used in Table 1 are shown. Magenta, green and white represent signals from XPP nuclear marker *PFA2 promoterHISTONE H2B-tdTomato, PIN7 promoter:PIN7-GFP* and cell wall stained with SR2200 respectively. Symbols shown below each image correspond to the cells on the respective sides of the image; the symbols on the left and right indicate data for the left and right cells, respectively. White arrows indicate anticlinal plasma membranes in which the GFP signal extended to the longitudinal side of the analyzed cells. Scale bar = 10 pm.

**Figure S9.**
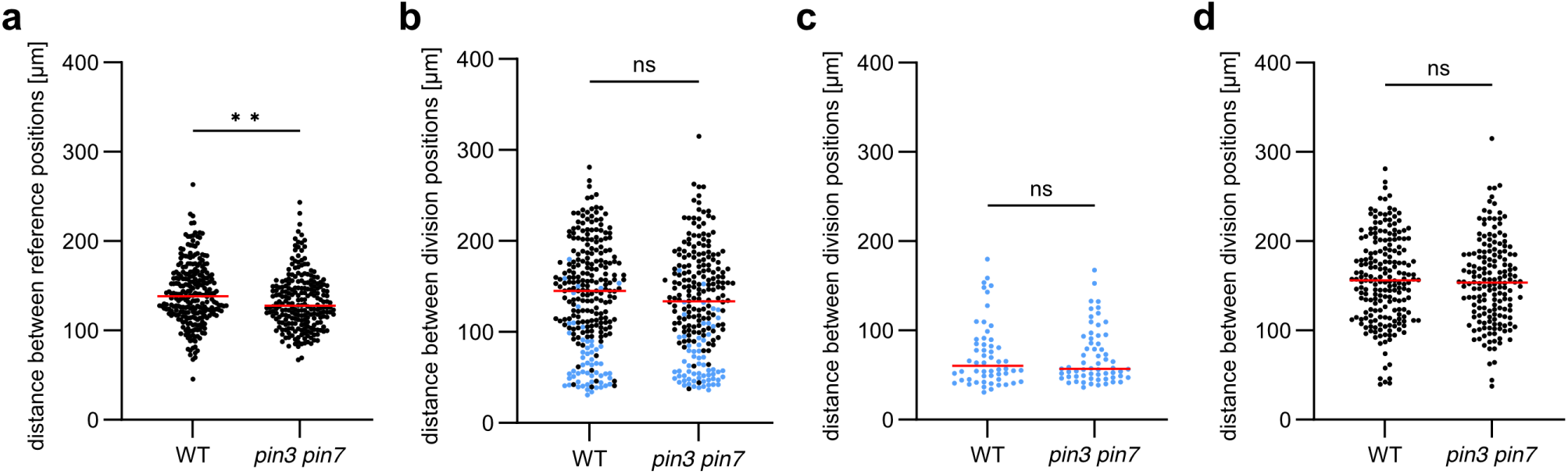
*pin3 pin7* mutants do not exhibit a significant defect in nuclear migration under 2 pM IAA treatment. 5-day-old WT and the *pin3 pin7* mutant seedlings expressing *PFA2 promoterHISTONE H2B-tdTomato* were treated with 2 pM IAA. Time-lapse imaging was performed with the method explained in Figure 3a. **a.** Distances between adjacent reference positions of all longitudinally adjacent XPP cells, n = 258 (WT), 232 *(pin3 pin7).* **b.** Distances between division positions of all longitudinally adjacent XPP nuclei. Blue dots represent distances between division positions of nuclei pairs with polar migration, n = 258 (WT), 232 *(pin3 pin7).* **c, d.** Blue and black dots from b were shown separately, n = 56 (WT), 62 *(pin3 pin7)* (c), n = 202 (WT), 170 *(pin3 pin7)* (d). Mann-Whitney test (**0.001 ≤ P < 0.01, *0.01 ≤ P < 0.05. ns, not significant (P ≥ 0.05)) was used to evaluate statistical significance (a-d).

**Figure S10.**
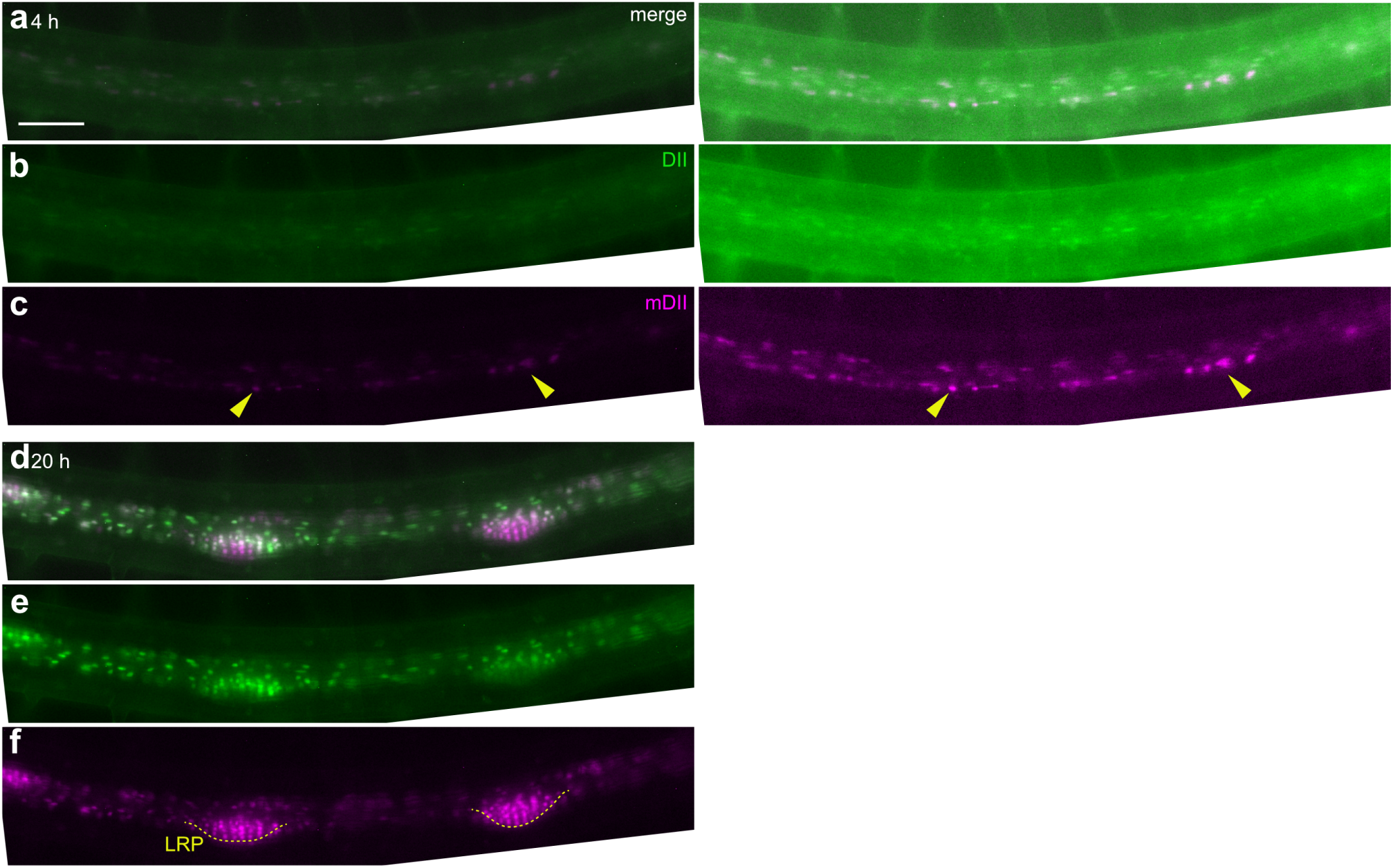
*RPS5A promoter:R2D2* signal patterns under 2 pM IAA treatment. Fluorescence microscopic time-lapse images of 5-day-old seedlings expressing original *R2D2* reporter *(RPS5A promoter:DII-n3xVenus/RPS5A promotenmDII-ntdTomato)* treated with 2 pM IAA. Merged images of DII-n3xVenus and mDII-ntdTomato (a, d), individual channel images of DII-n3xVenus (b, e) and mDII-ntdTomato (c, f) at 4 hours(a-c) or 20 hours (d-f) of IAA treatment are shown. The right images in a-c show the same images as those on the left, with adjusted contrast for clarity. Yellow arrowheads in d indicate the positions where asymmetric cell divisions occur, leading to the formation of dome-shaped LRPs (yellow dashed lines in f). Scale bar = 100 pm.

